# Effects of prolonged fixation on vascular biomarkers in postmortem human brains

**DOI:** 10.64898/2026.02.27.708588

**Authors:** Eve-Marie Frigon, Weiya Ma, Cecilia Tremblay, Dominique Mirault, Gustavo Turecki, Naguib Mechawar, Denis Boire, Josefina Maranzano, Mahsa Dadar, Yashar Zeighami

## Abstract

Postmortem human brains stored in brain banks are important research resources to study the mechanisms underlying normal brain functions as well as various neurodegenerative disorders. Immunohistochemical (IHC) and histochemical (HC) staining have been used to examine human brains fixed in neutral-buffered formalin (NBF) for months, years, and even decades. As such, it is essential to establish the effects of prolonged fixation in NBF on both IHC and HC stains. Previously, we found that prolonged NBF fixation resulted in differential effects on IHC and HC staining on postmortem brains. In this study, we further examined the effects of prolonged fixation on IHC stains targeting 6 antigens and 2 HC stains of known biomarkers of cerebrovascular diseases in prefrontal cortex of human brains fixed for 1, 5, 10, 15, and 20 years. The IHC targets included microvasculature markers of the blood brain barrier (Collagen-IV and Claudin-5), a type III intermediate filament marker (Vimentin), an activated microglia marker (CD68), a biomarker for oligodendrocytic myelin proteolipid protein (PLP) and a marker for iron accumulation (Ferritin). The HC included Masson’s Trichrome Stain (MTS) and Bielschowsky silver stain (BSS). We found that staining intensities of Ferritin, Vimentin, Collagen-IV and BSS decreased with prolonged fixation, while no significant differences were observed in the staining intensity of other markers. Hence, these differential alterations should be taken into consideration when interpreting the results from processed tissues with prolonged fixation. We recommend performing IHC and HC staining for human brains with the same fixation times to offset any impact on downstream neuropathological analyses, as well as adding the fixation duration as a covariate in the analysis.

## 1. Introduction

Emerging evidence suggests that the development of neurodegenerative diseases (NDD), including Alzheimer’s disease (AD), may be preceded and exacerbated by vascular changes. Cerebrovascular disease (CVD) has been identified as an important contributor to age-related dementias, frequently co-existing alongside neurodegenerative pathologies and impacting disease onset and progression rate [1–3]. Markers of CVD, namely white matter hyperintensities (WMH), infarcts, and microbleeds, can be detected using structural magnetic resonance imaging (MRI) [4, 5]. However, structural MRI lacks the sensitivity and specificity required to reliably characterize the underlying microscopic pathologies associated with CVD. These imaging findings can instead be validated through postmortem histological evaluation of the affected tissues, which remains the gold standard for identifying the cellular and molecular alterations associated with CVD lesions [6–8].

Histology procedures such as immunohistochemistry (IHC) and histochemistry (HC) are commonly used for cellular assessments (i.e. number, location, or morphology) of cerebral tissues, confirming the presence and severity of various diseases-related pathological processes [9–11]. IHC and HC staining quality highly depends on tissue fixation quality [12, 13] and can therefore be affected by differences in fixative chemicals used [14, 15], fixation length [16] or postmortem interval (PMI) [17]. Since postmortem tissue requests are typically submitted to brain banks after sufficient numbers of cases meeting specific inclusion criteria have accumulated, relatively few studies are able to consistently analyze samples shortly after donation. Consequently, most tissues are fixed in 10% neutral buffered formalin (NBF) for months, years, or even decades before being used for research [18, 19], which results in substantial variability in fixation duration both within and between studies, highlighting the need for a thorough assessment of its impact on downstream results.

NBF preserves tissue integrity by hardening the tissue to prevent decomposition, degradation, and shrinkage [18, 20, 21]. However, it also induces crosslinking of proteins and nucleic acids within tissues [22, 23], which in turn, can mask antigenic epitopes and reduce immunoreactivity in IHC [13, 24]. To circumvent this drawback, antigen retrieval approaches are widely used to reverse the NBF-induced crosslinking in human brain tissues with prolonged fixation before IHC staining [25–28]. Prior studies have shown that the effect of prolonged fixation on IHC and/or HC staining in human brains fixed for periods ranging from weeks to years can be either negative or unaltered [29–32]. A recent study from our team showed that longer fixation times were associated with reduced staining intensity of the neuronal nuclei marker NeuN, and the ionized calcium-binding adapter molecule 1 (Iba1) as a microglia marker, while other markers (i.e., glial fibrillary acidic protein (GFAP)) and HC staining (i.e., Hematoxylin and Eosin (H&E)) intensity remained stable across fixation durations [16]. Furthermore, it has been previously suggested that gray matter (GM) and white matter (WM) regions could be differentially affected by the diffusion of chemical fixatives, as shown using MRI [33], however, this effect on HC and IHC studies has not been investigated. Taken together, these data suggest that prolonged fixation can produce differential effects on the quality of histological staining and neuropathological quantification and assessment. Since IHC and HC staining can be influenced by formaldehyde fixation but are necessary for identification of neuropathology and underlying mechanisms in CVD, it is essential to determine whether prolonged fixation alters the expression and detectability of biomarkers widely used to diagnose and study CVD.

Therefore, the present study aimed to determine whether prolonged fixation in NBF affects the staining intensity of antigens related to the microvasculature, the blood-brain barrier (BBB), and other CVD or NDD biomarkers in postmortem human brains, specifically within WM and GM regions.

## 2. Materials and methods

### 2.1 Postmortem human brain samples

Brain samples were obtained from the Douglas Brain Bank (DBB; https://douglasbrainbank.ca). Consistent with the DBB protocol, hemispheres were separated by a sagittal cut in the middle of the brain, brainstem, and cerebellum upon receipt [34]. One unsliced hemisphere (right or left, in alternation) was fixed in 10% NBF. For this study, 20 prefrontal cortex (PFC) blocks from specimens fixed for 1, 5, 10, 15, and 20 years (n=4 per group, in total, n=20) were requested. All specimens were consistently processed and fixed in the same solution (i.e. 10% NBF), with the sole difference being the duration of fixation.

All brains were recruited from the DBB after obtaining consent by next-of-kin in collaboration with the Quebec Coroner’s Office. All experimental procedures were approved by Human Research Ethics Committees of McGill University and Douglas Mental Health University Institute. The cases were recruited from the suicide brain bank, in which individuals with evidence of drugs or psychotropic medications or documented history of neurological disorders or head injury were excluded from the study. However, since no brain tissue was available for the 10-years group in that cohort, cases for this group were recruited from the aging and dementia brain bank. These brains have a full neuropathological evaluation, and their neuropathological diagnosis and presence of CVD are reported in Table 1. Table 1 also presents the characteristics of the samples included in this study.

**Table 1.**
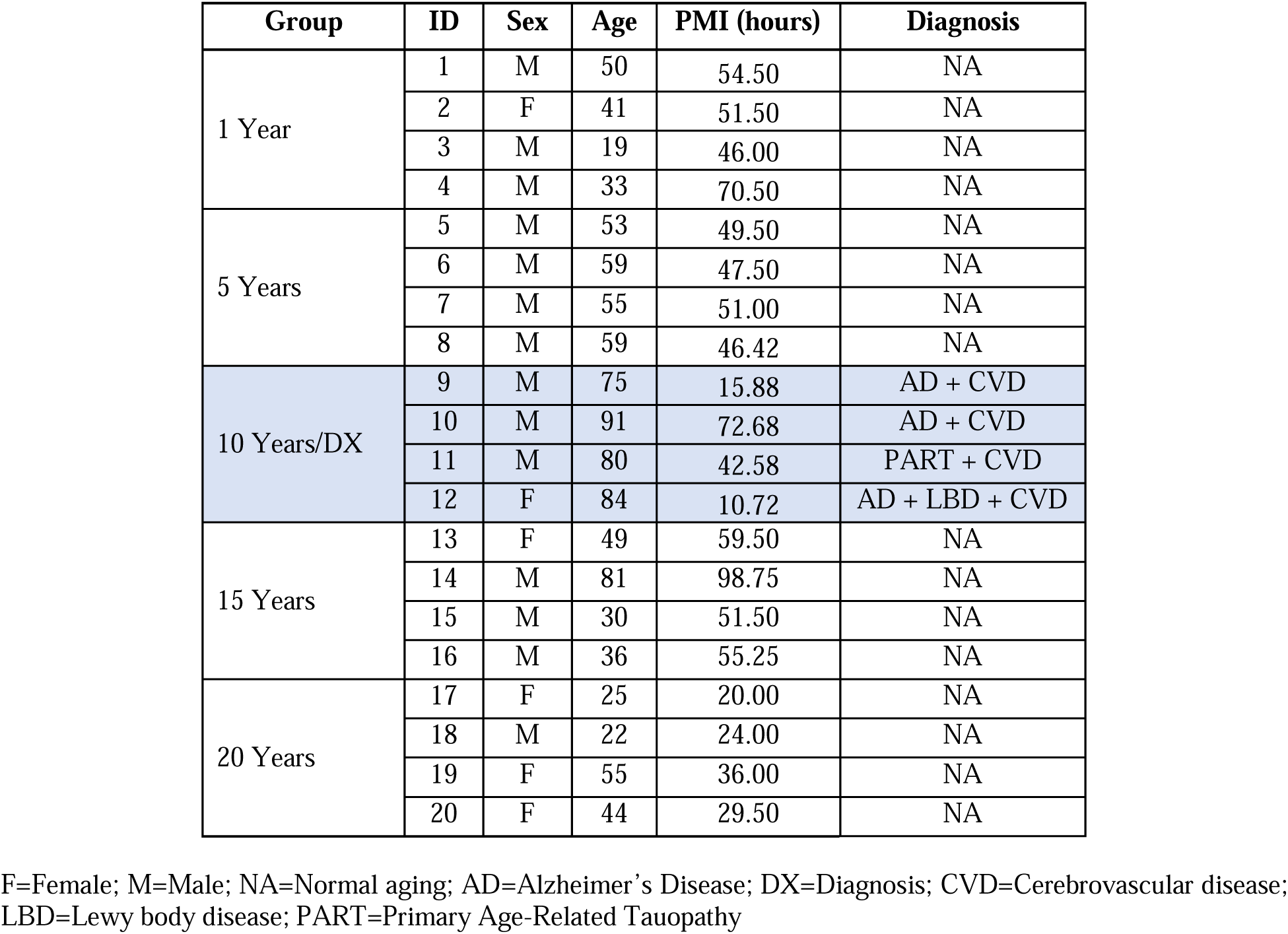
Demographics of the specimens used in this study.

### 2.2 Cryostat sectioning of PFC samples

Upon reception, all blocks were transferred to 30% sucrose in 0.1M phosphate buffered 0.9% saline (PBS) (pH 7.4) for cryoprotection until cryostat sectioning. PFC blocks were embedded in M1 embedding medium (Fisher Scientific, Saint-Laurent, Quebec, Canada) and cut into sections of either 50-μm (for IHC, see section 2.3) or 20-μm thickness (for HC, see section 2.4) (Leica, Feasterville, Pennsylvania, USA). The 20-µm-thick sections were cut from the central portion of the blocks, sequentially mounted on gelatin pre-coated slides, and stored at −80oC until use. The 50-µm-thick sections were sequentially collected in cryoprotectant contained in 24-well culture plates and stored at −20oC until used for IHC staining.

### 2.3 Immunohistochemistry

Table 2 presents the antigens that were used to assess microvasculature and elements of the BBB structure (collagen-IV, vimentin, and claudin-5), neuroinflammation marker of activated microglia using cluster of differentiation 68 (CD68), oligodendrocytic myelin proteolipid protein (PLP), and brain iron accumulation using anti-ferritin.

**Table 2.**
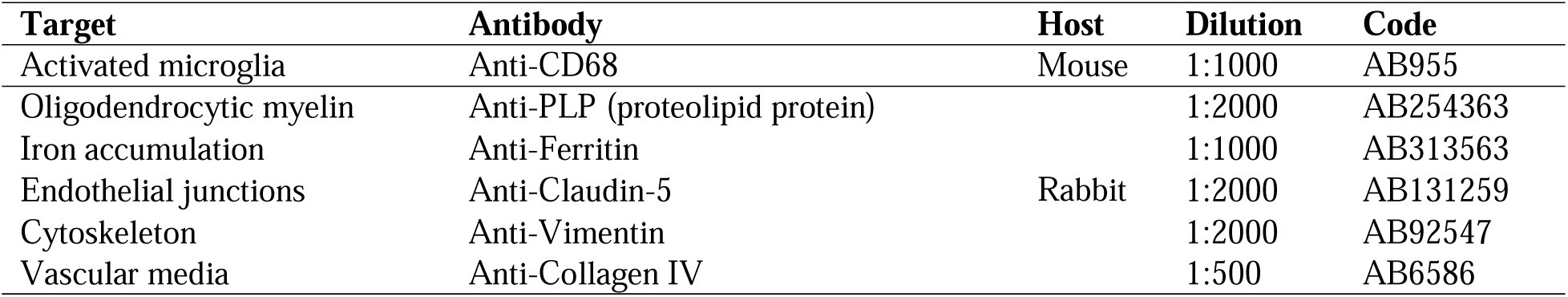
Summary of primary antisera used in this study.

For free-floating immunostaining, the 50 μm-thick PFC sections were first rinsed in PBS, then transferred into Eppendorf tubes with antigen retrieval buffer (10mM sodium citrate, 0.05% Tween 20, pH 6.0). The tubes were heated to 105°C inside the slots of a hot plate for 25 min. Then the tubes were cooled down on wet ice, and sections were rinsed with cold double-distilled water (_dd_H_2_O) for 10 min. Sections were transferred into PBS containing 0.05% Triton-100 (PBS+T) in wells of a 24-well culture plate, quenched in 0.03% H_2_O_2_ for 15 min to eliminate the endogenous peroxidase in the tissue, and then incubated in 50% ethanol for 1 hour to increase antibody penetration into the tissues. Sections were incubated in 10% normal goat serum (NGS) and 1% bovine serum albumin diluted in PBS+T for 1 to 2 hours to block the endogenous non-specific antigenic sites. Finally, sections were incubated at room temperature in one of the six monoclonal anti-human antibodies (either hosted in mouse or rabbit), as presented in Table 2. Sections were next incubated in a biotinylated goat anti-mouse or in a biotinylated goat anti-rabbit IgG (1:200, MJS Biolynx Inc., Brockville, Ontario, Canada) for 1 hour and further processed by using an Elite ABC kit (MJS Biolynx Inc.) for another hour according to the manufacturer’s instructions. Finally, the immunoprecipitates were developed using 3,3-diaminobenzidine (DAB, Sigma-Aldrich, Oakville, Ontario, Canada) as the chromogen and enhanced by the glucose oxidase-nickel-DAB method [35]. Between incubations, sections were washed thoroughly in PBS+T twice. Finally, sections were mounted on the gelatin-coated slides (Fisher Scientific), dehydrated in ascending ethanol solutions, cleared in xylene, and cover-slipped with xylene-based mounting medium (Micromount, Leica).

### 2.4 Histochemistry

The 20µm-thick sections mounted on gelatin-coated slides were removed from the −80°C freezer and air-dried. For dehydration, section-mounted slides were treated sequentially in xylene (20 min x 2), 100% ethanol (2 min x 2), 95% ethanol (2 min x 2), and 70% ethanol (2 min x 2).

We first used the Masson’s Trichrome Stain (MTS) Kit (Abcam, Cat Ab150686), where slides were immersed in pre-heated Bouin’s fluid (60 °C) for 60 min followed by a 10 min cooling period. Slides were then rinsed under tap water until sections were completely clear. After briefly rinsing in _dd_H_2_O, slides were stained in the working Weigert’s iron hematoxylin solution for 5 min before rinsing under tap water for 2 min. Then, slides were immersed in Biebrich Scarlet/Acid Fuchsin Solution for 15 min before rinsing briefly in _dd_H_2_O. Slides were differentiated in phosphomolybdic/phosphotungstic acid solution for 12 min or until slides were no longer red. Aniline Blue Solution was then applied to slides for 10 min before rinsing in _dd_H_2_O. Slides were finally immersed in 1% acetic acid solution for 5 min before being dehydrated in ascending ethanol and cleared in xylene.

For Bielschowsky silver staining (BSS), slides were immersed in 10% silver nitrate solution (Sigma-Aldrich) in the 45 °C oven for 20 min. Slides were rinsed thoroughly with _dd_H_2_O (3x). In the same silver nitrate solution, concentrated ammonium hydroxide (Fisher Scientific) was added drop by drop while mixing with a stirring bar until the solution was completely clear. Slides were immersed in the working ammoniacal silver solution for 20 min (45 °C). Next, slides were immersed in the working developer solution (37-40% formaldehyde, citric acid, nitric acid, all from Sigma-Aldrich) for 1 min and in 1% Ammonium hydroxide (Fisher Scientific) for 1 min. After rinsing in _dd_H_2_O (3x), slides were immersed in 5% sodium thiosulfate (Sigma-Aldrich) for 5 min. After rinsing in _dd_H_2_O (3x), slides were dehydrated in 95% ethanol (2 min x 2) and 100% ethanol (2 min x 2), and cleared in xylene (5 min x 2) before being cover-slipped.

### 2.5 Image capture, quantification, and statistical analysis

Ten brightfield images (20X) per specimen were acquired in five regions of interest (ROIs) randomly selected within the WM and GM. With four specimens per group, this resulted in 40 images per group of comparison (i.e. a total of 200 images per staining type). Image acquisition parameters were kept consistent between ROIs, specimens, and groups of comparison; images were captured at the same magnification (UPlanSApo 20X/0.75∞0.17/FN26.5objective), with the same threshold of illumination or exposure time on a brightfield Olympus microscope (BX51W1) controlled by Neurolucida software (MBF Bioscience). Each captured field resulted in an image with a dimension of 2752 x 2192 pixels (623.44 x 499.26 μm) using a DV-HR-CLR Lumina High Resolution Color Camera 1”CCD Sensor. For quantification of Vimentin, CD68, Claudin-5, and Collagen-IV staining intensity, each image was measured by automatic extraction of threshold-based intensity from the 8-bit images using ImageJ (ImageJ ver.1.53, Softonic, Barcelona, Spain). PLP staining images were converted to 8-bit images prior to extracting their mean gray values using an ImageJ macro plug-in. Ferritin, MTS and BSS stains were assessed by extracting their mean RGB values using an in-house MATLAB script.

The mean staining intensity values for GM and WM ROIs of all fixation groups were statistically compared using linear mixed-effect models to account for within- and between-sample variability given the repeated measures obtained from each sample. The models included specimen ID as a categorical random variable as well as sex, age and PMI as covariates:

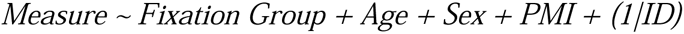

*Measure* indicates the GM or WM staining intensity values for different stains of interest, and *Fixation Group* indicates a categorical variable contrasting prolonged fixation groups (i.e. 5 years, 10 years, 15 years, and 20 years) against the 1-year group. The results were corrected for multiple comparisons using the False Discovery Rate (FDR) control procedure, with a significance threshold of 0.05 [36].

## 3. Results

Mean age at time of death for the 1-, 5-, 10-, 15- and 20-years groups were 35.7 (range 19-50), 56.5 (range 53-59), 82.5 (range 75-91), 49 (range 30-81) and 36.5 (range 22-55) years, respectively. The specimens included in the 10-year group were significantly older than all other groups (p<0.05) and had postmortem evidence of CVD and NDD (Table 1). Male to female ratios were 3:1, 4:0, 3:1, 3:1 and 1:3, respectively. PMIs were not significantly different across groups.

### 3.1 Immunohistochemical staining

Table 3 summarizes the results of the mixed effect models used to assess the differences between mean staining intensities of the prolonged fixation groups (5-years, 10-years, 15-years and 20-years) against the 1-year group, adjusted for age, sex, and PMI. Presented values reflect the t statistics of contrasting the groups of interest against the reference 1-year group, and stars indicate the significance of the results following FDR correction.

**Table 3.**
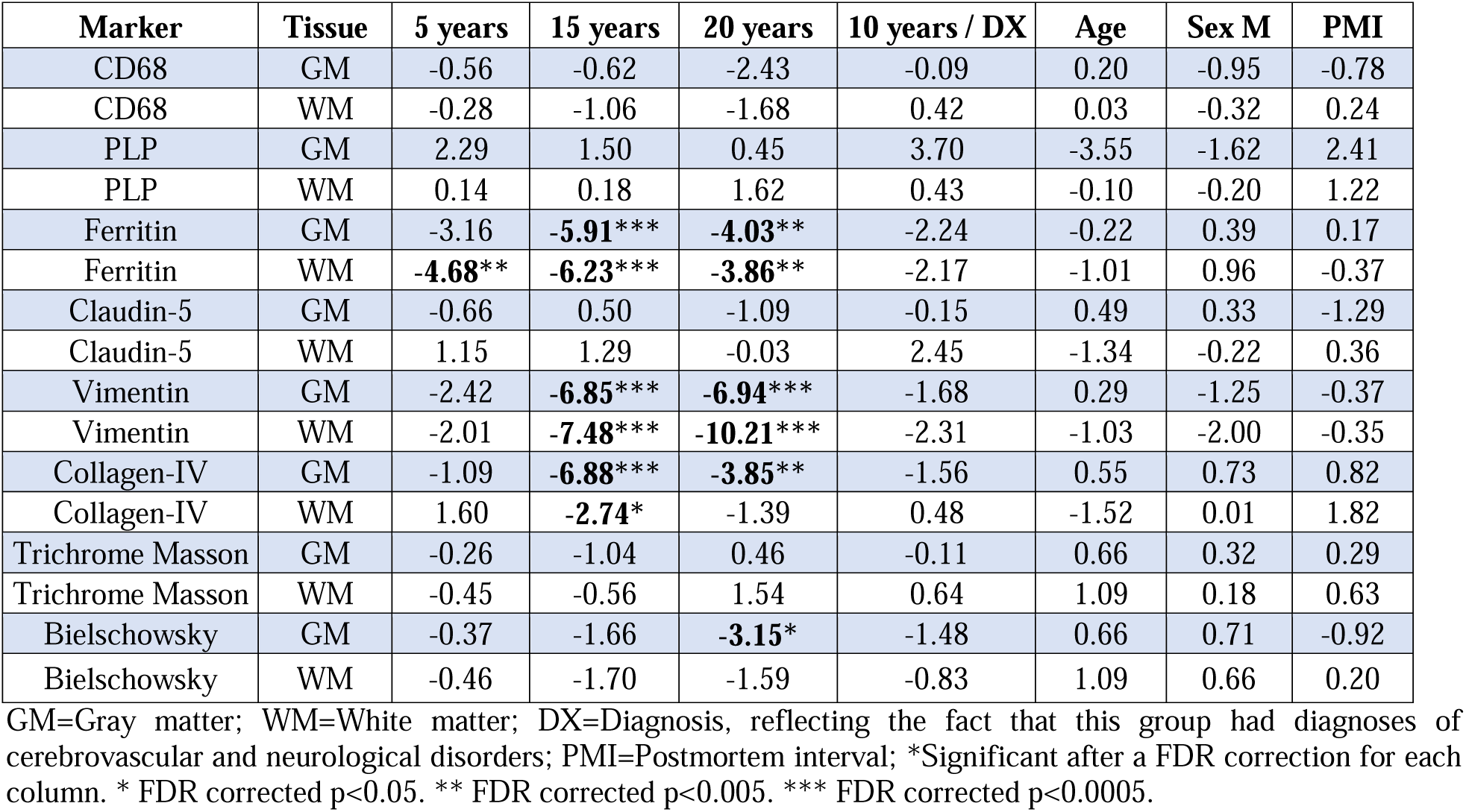
Differences in mean staining intensities of the prolonged fixed groups compared against the 1-year group.

CD68 labeled activated microglia in the 5 groups (Figure 1). In the GM and WM regions, the staining intensities decreased with fixation time and were higher in the 10 Year/DX group, although these differences were not statistically significant (Figure 1A-B, Table 3).

**Figure 1.**
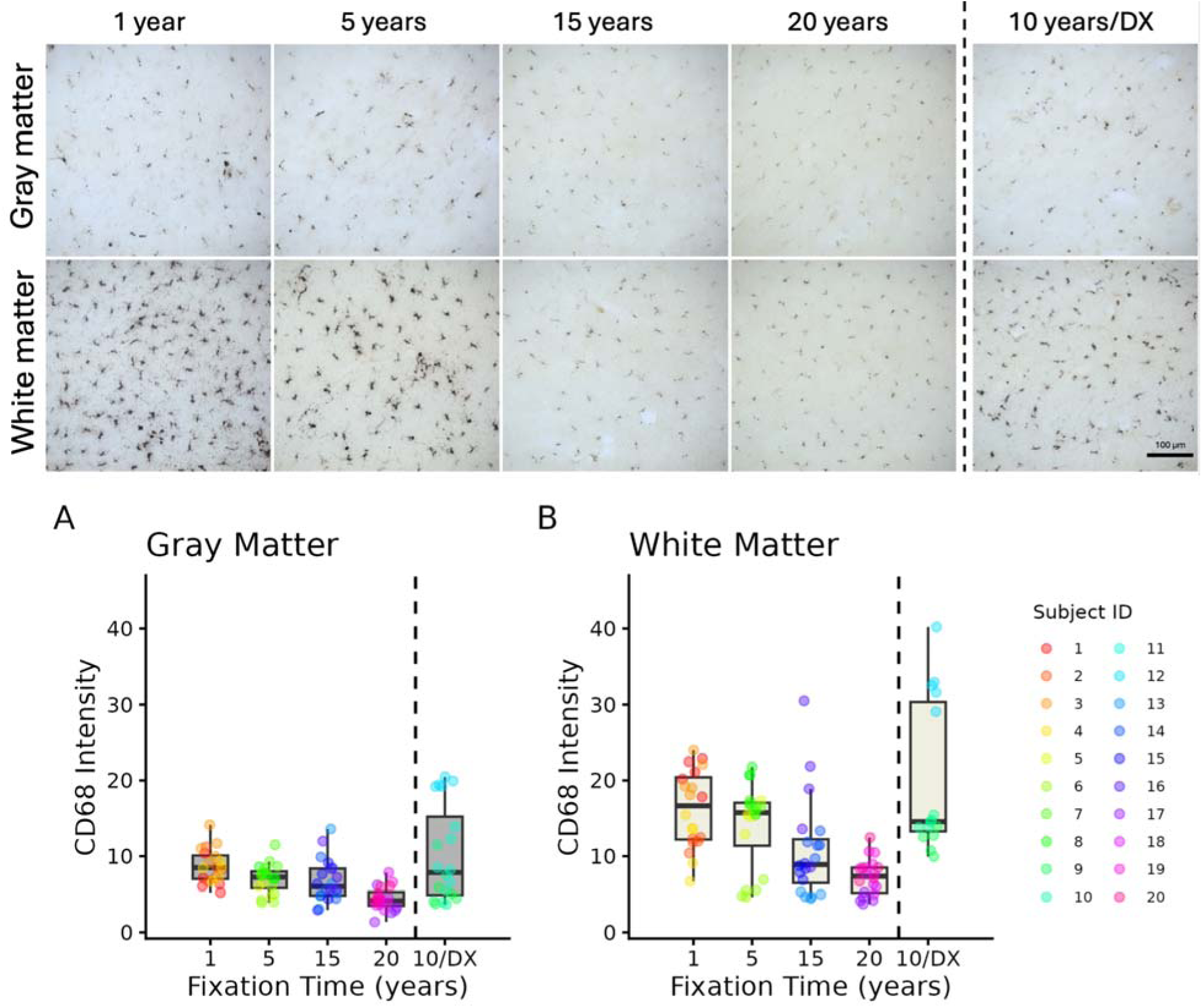
Distribution of CD68-immunoreactive activated microglia profiles in the gray and white matter (20X) of the PFC sections of human brains fixed for 1, 5, 10, 15, and 20 years. A) Mean staining intensity of the gray matter in the five groups. B) Mean staining intensity of the white matter in the five groups. Mean staining intensity was measured using a threshold-based intensity in ImageJ. DX=Diagnosis; ID=Identification.

PLP stained myelin fibers in the GM and WM of the 5 groups (Figure 2). In GM regions, the gray values were not different across the five fixation groups (Figure 2A, Table 3). WM staining intensity of PLP had an increasing trend with prolonged fixation (Figure 2B), indicating that the fibers were less differentiated from the background (i.e., lower staining quality), but this was not statistically significant (Table 3).

**Figure 2.**
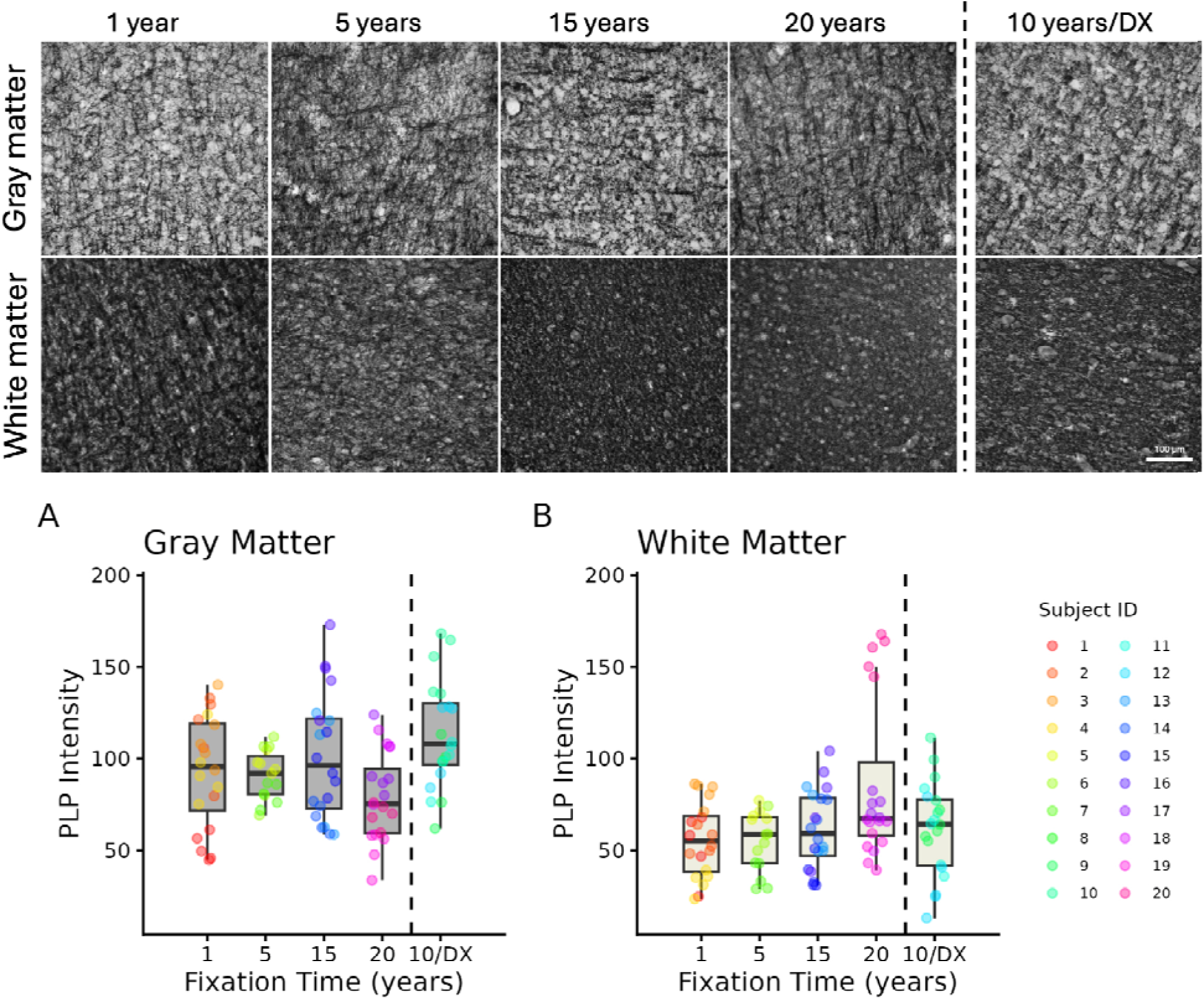
Distribution of PLP-immunoreactive myelin fibers profiles in the gray and white matter (20X) of the PFC sections of human brains fixed for 1, 5, 10, 15, and 20 years. A) Mean staining intensity of the gray matter in the five groups. B) Mean staining intensity of the white matter in the five groups. Mean staining intensity was measured using a macro plugin in ImageJ. DX=Diagnosis; ID=Identification.

Ferritin marker labeled dark deposits of iron and microglia-shaped cells in the five groups (Figure 3). In the GM, the 1-year group staining intensity (median=139.85) was significantly higher than in the 5-years (median=59.09, p<0.01), 10-years (median=48.30, p<0.05), 15-year (median=38.60, p<0.001) and 20-years (median=48.00, p<0.01) groups (Figure 3A). However, the FDR corrected results in the 5-years did not reach the significance threshold (Table 3). In WM regions, the 1-year group staining intensity (median=68.71) was significantly higher than in the 5-years (median=13.75, p<0.001), 10-years (median=23.50, p<0.05), 15-years (median=9.31, p<0.001) and 20-years (median=23.83, p<0.01) groups (Figure 3B, Table 3).

**Figure 3.**
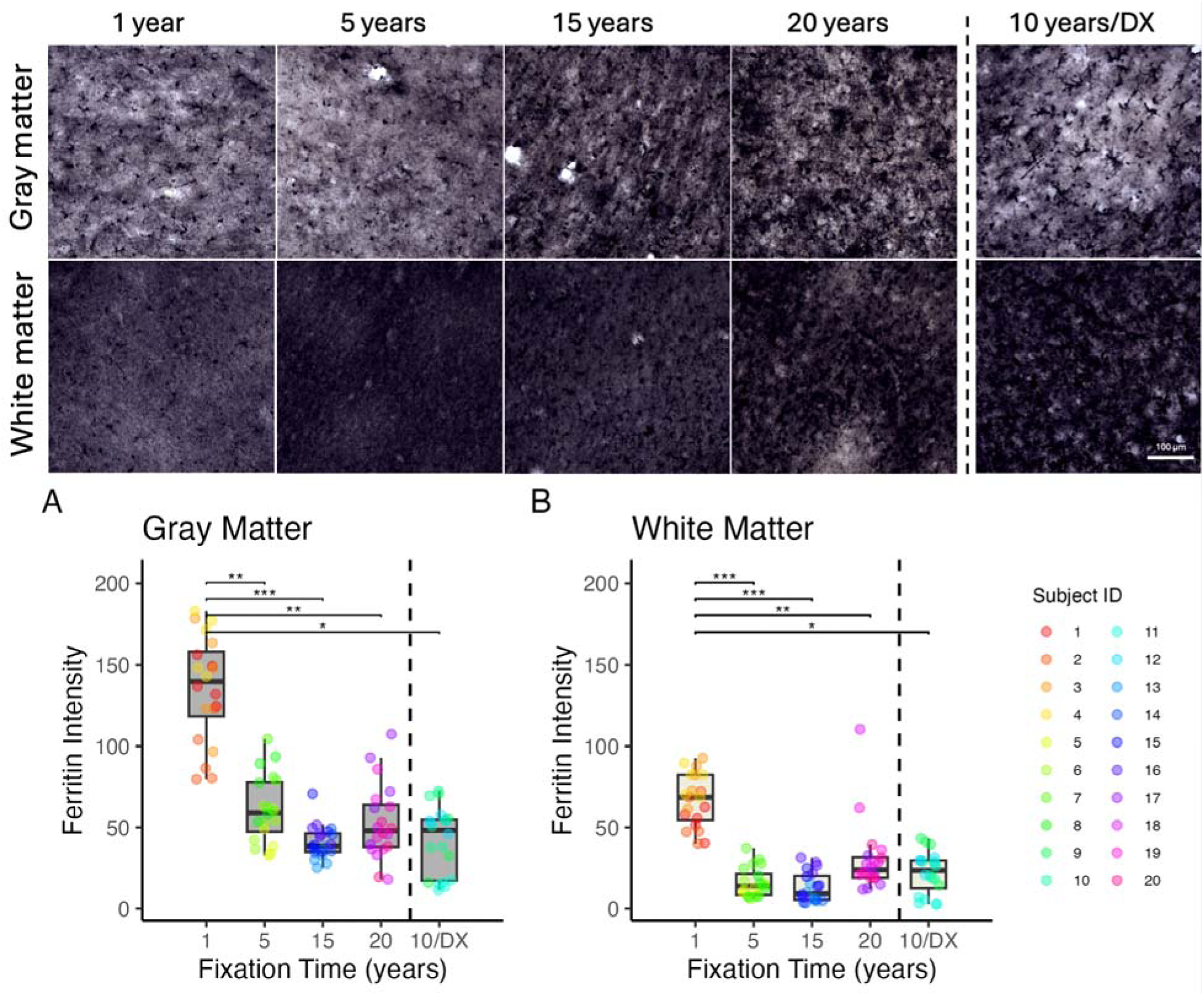
Distribution of Ferritin-immunoreactive activated microglia and iron deposits profiles in the gray and white matter (20X) of the PFC sections of human brains fixed for 1, 5, 10, 15, and 20 years. A) Mean staining intensity of the gray matter in the five groups. B) Mean staining intensity of the white matter in the five groups. Mean staining intensity was measured based on RGB intensity values. *uncorrected p<0.05. **uncorrected p<0.005. ***uncorrected p<0.0005. DX=Diagnosis; ID=Identification.

Figure 4 shows blood vessels positively stained for Claudin-5 in GM and WM regions of the five fixation groups. We found no significant differences in the staining intensity of the GM ROIs (Figure 4A). The 10-years group showed higher staining intensities (median=12.19) compared to the 1-year group (median=7.44; p<0.05) in WM regions (Figure 4B), but this did not retain significance after a FDR correction (Table 3).

**Figure 4.**
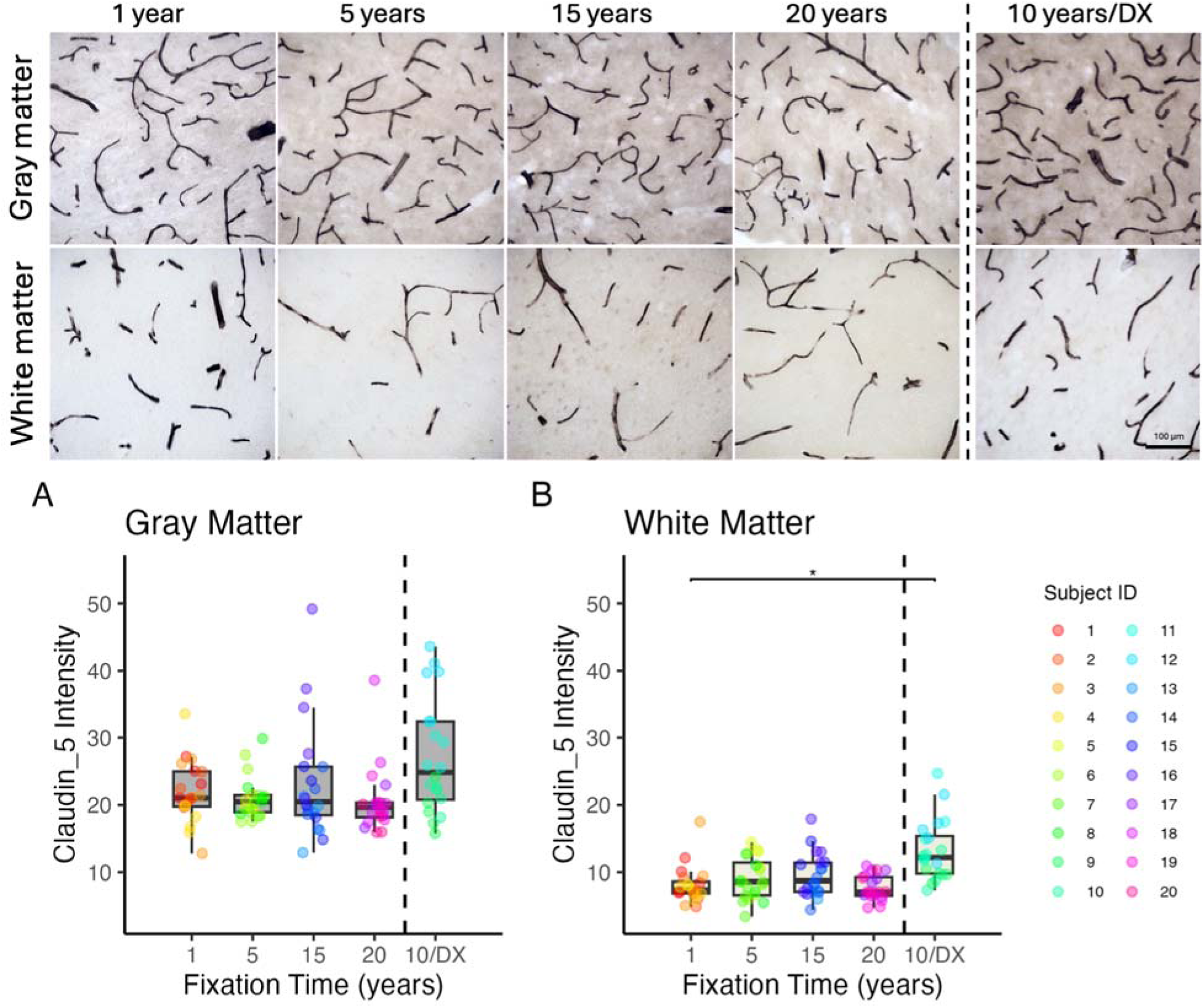
Distribution of Claudin-5-immunoreactive profiles in the gray and white matter (20X) of the PFC sections of human brains fixed for 1, 5, 10, 15, and 20 years. A) Mean staining intensity of the gray matter in the five groups. B) Mean staining intensity of the white matter in the five groups. Mean staining intensity was measured using a threshold-based intensity in ImageJ. *uncorrected p<0.05. DX=Diagnosis; ID=Identification.

Vimentin marker stained blood vessels in the first four groups, while staining was completely absent in the 20-years group. Vimentin also labeled astrocytes-shaped cells in the 1-year group (Figure 5). In the GM, we found that the 1-year group (median=31.92) showed significantly higher staining intensity than the 5-years (median=21.18; p<0.05), 15-years (median=3.82; p<0.001) and 20-years (median=0; p<0.001) groups (Figure 5A). However, the difference with the 5-years group did not retain significance after a FDR correction (Table 3). In the WM ROIs, median staining intensity values of the 1-year group (median=17.66) were significantly higher than in the 15-years (median=4.07; p<0.001), 20-years (median=0; p<0.001) and 10-year (median=7.08; p<005) groups (Figure 5B). However, the 10-years group did not retain significance after a FDR correction (Table 3).

**Figure 5.**
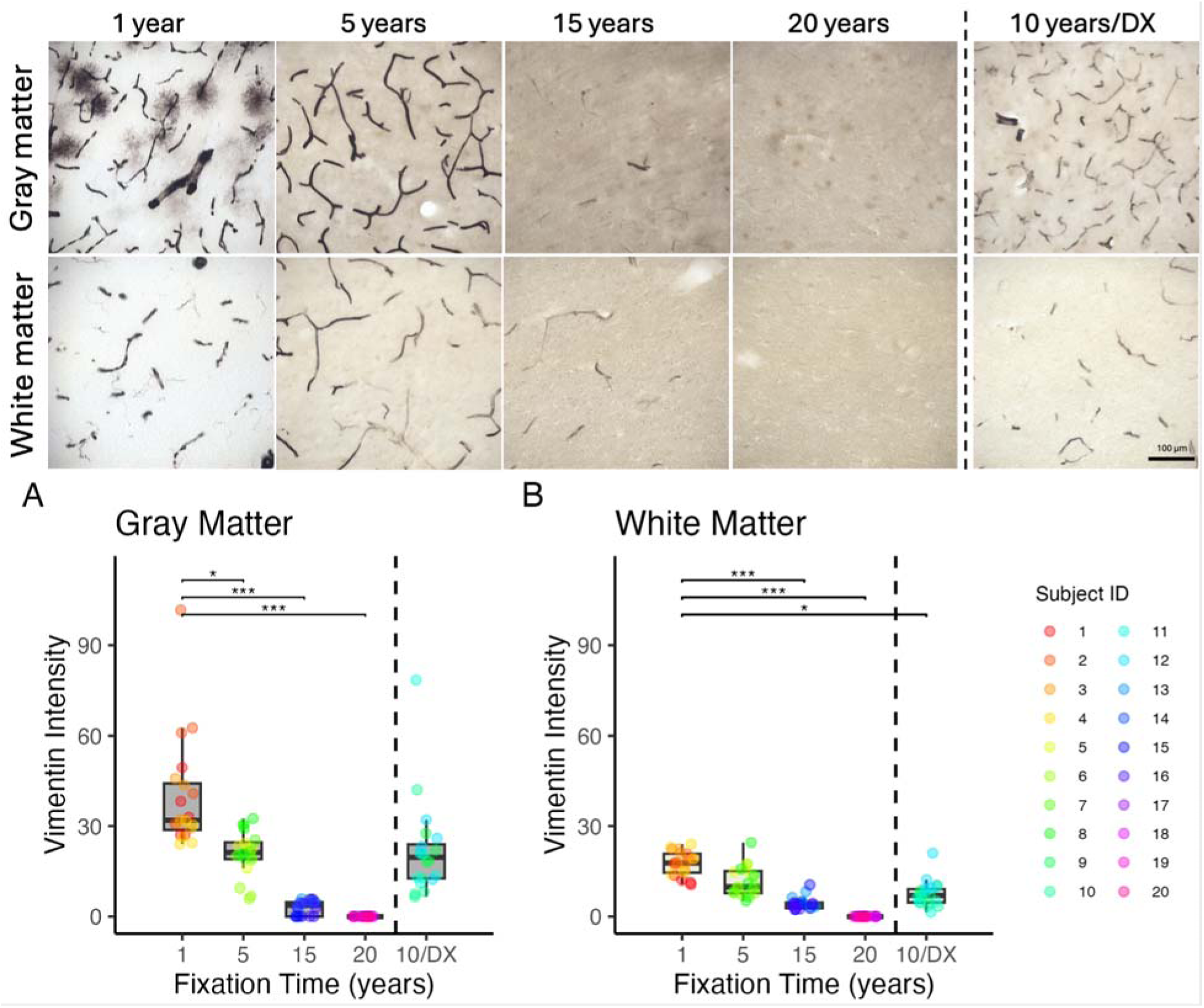
Distribution of Vimentin-immunoreactive profiles in the gray and white matter (20X) of the PFC sections of human brains fixed for 1, 5, 10, 15, and 20 years. A) Mean staining intensity of the gray matter in the five groups. B) Mean staining intensity of the white matter in the five groups. Mean staining intensity was measured using a threshold-based intensity in ImageJ. *uncorrected p<0.05. ***uncorrected p<0.0005. DX=Diagnosis; ID=Identification.

Collagen-IV stained blood vessels as well as glial cell bodies in the five groups (Figure 6). In the GM, staining intensity of the 1-year (median=39.78) group was significantly higher than in the 15-years (median=6.03; p<0.001) and 20-years (median=10.37; p<0.01) groups (Figure 6A, Table 3). In the WM, staining intensity of the 1-year (median=19.09) group was significantly higher than in the 15-years group (median=2.15; p<0.001) (Figure 6B, Table 3).

**Figure 6.**
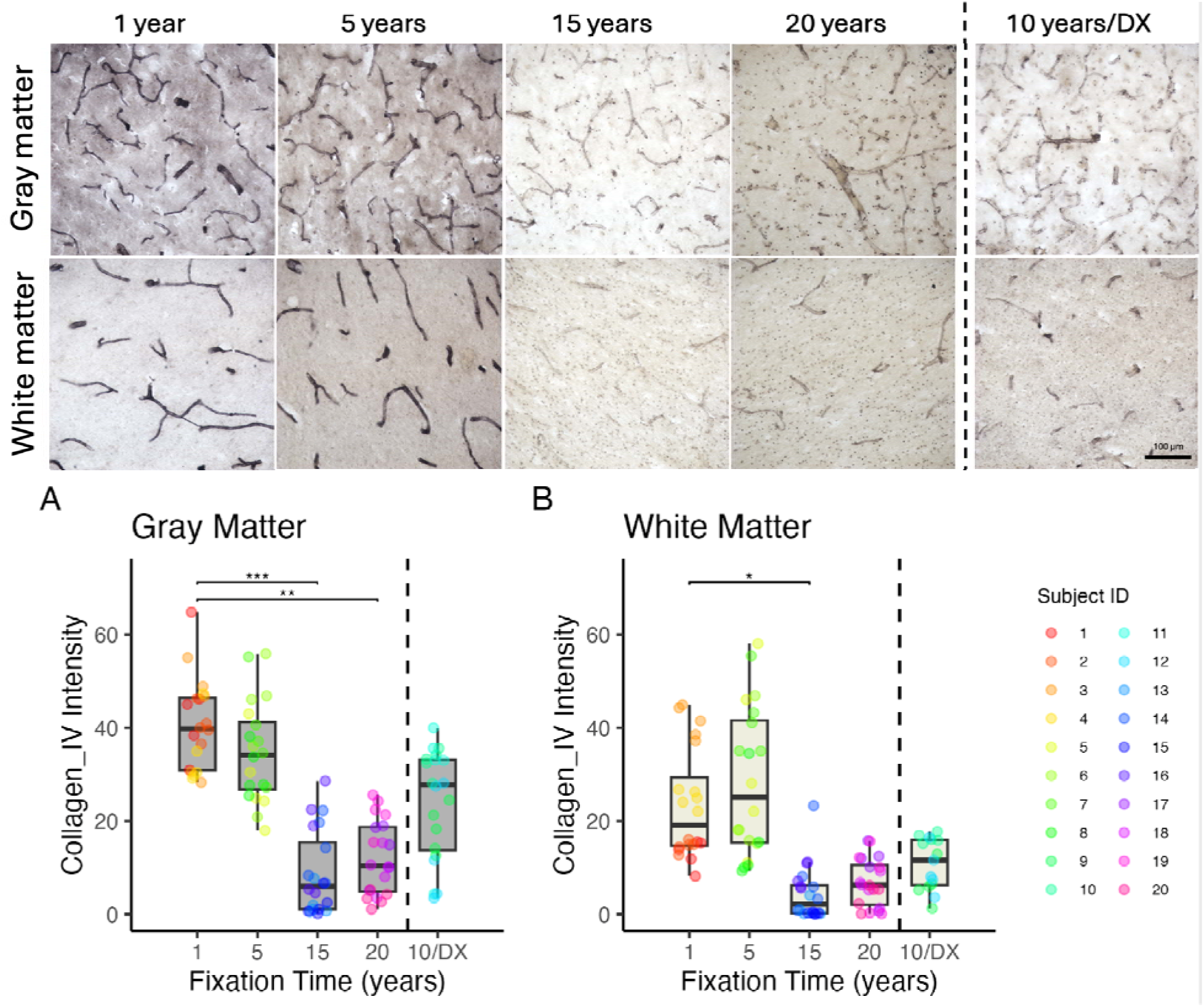
Distribution of Collagen-IV-immunoreactive profiles in the gray and white matter (20X) of the PFC sections of human brains fixed for 1, 5, 10, 15, and 20 years. A) Mean staining intensity of the gray matter in the five groups. B) Mean staining intensity of the white matter in the five groups. Mean staining intensity was measured using a threshold-based intensity in ImageJ. *uncorrected p<0.05. **uncorrected p<0.005. ***uncorrected p<0.0005. DX=Diagnosis; ID=Identification.

### 3.2 Histochemical staining

#### 3.2.1 Masson’s trichrome staining

MTS stained neuropils in purple, cell nuclei in dark blue, and blood cells in red (Figure 7). We found no significant differences in the staining intensity of the GM nor the WM regions in brain sections fixed for 1 year, 5 years, 10 years, 15 years or 20 years (Figure 7A, 7B, Table 3). However, we observed a trend where the increased staining intensity with prolonged fixation or in brain specimens with a NDD diagnosis, reflecting a lower differentiation between the labeled cells and fibers and the background (i.e., lower staining quality).

**Figure 7.**
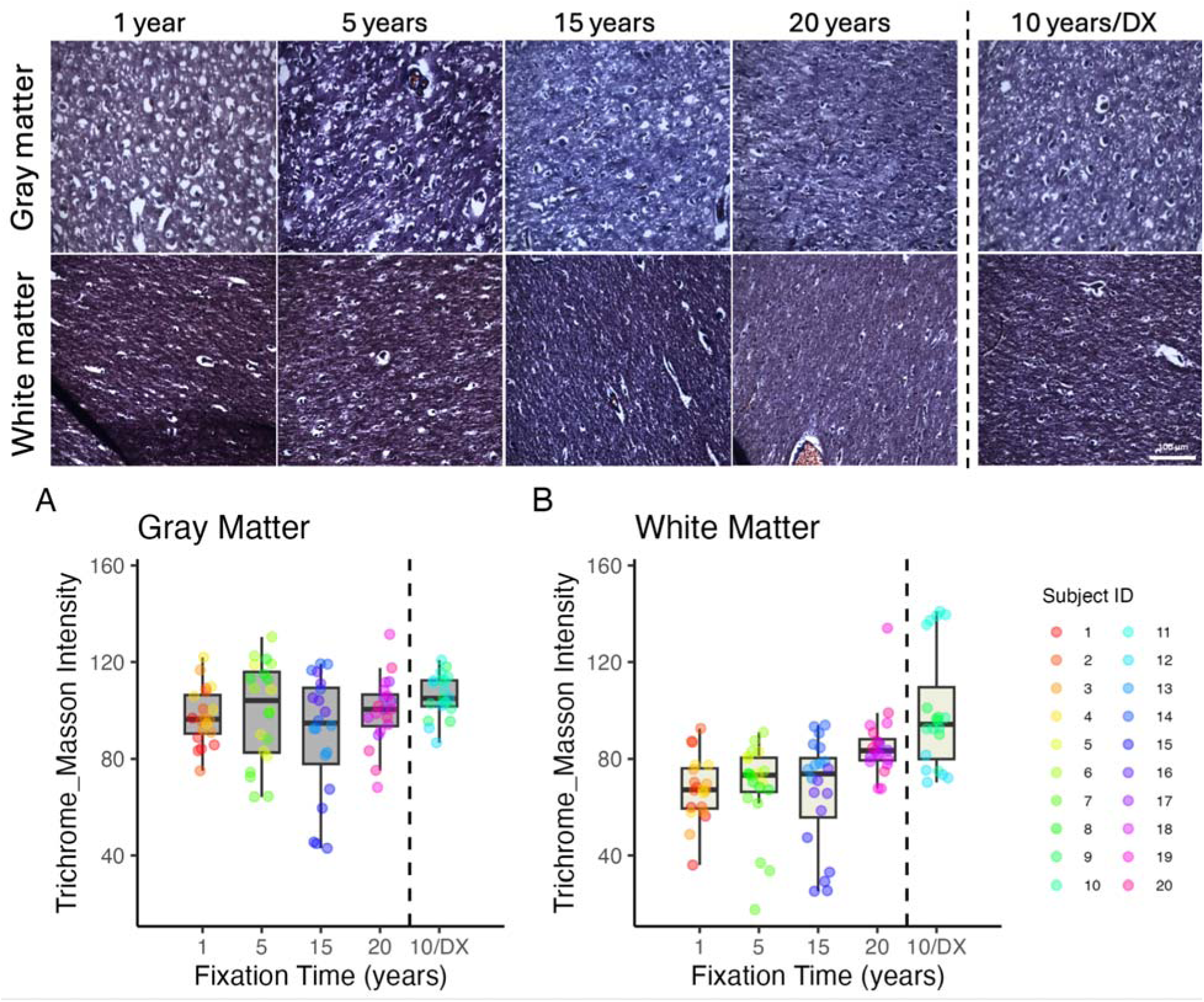
Masson’s Trichrome staining in the gray matter and white matter (20X) of the PFC sections of human brains fixed for 1, 5, 10, 15, and 20 years. A) Mean staining intensity of the gray matter in the five groups. B) Mean staining intensity of the white matter in the five groups. Mean staining intensity was measured based on RGB intensity values. DX=Diagnosis; ID=Identification.

#### 3.2.2 Bielschowsky silver staining

BSS labeled axons and fibers throughout the GM and WM regions, where thick axonal fiber bundles were stained brown while the background of neuropils was stained yellow (Figure 8). Three out of four cases of the 10-years group presented high levels of AD neuropathologic changes, senile plaques, and neurofibrillary tangles in the GM when stained with BSS. In the GM, the 1-year (median=79.23) showed significantly higher RGB values than in the 20-year group (median=46.88; p<0.01) (Figure 8A, Table 3). We found no significant difference in th WM ROIs in the BSS staining intensity with prolonged fixation (Figure 8B, Table 3).

**Figure 8.**
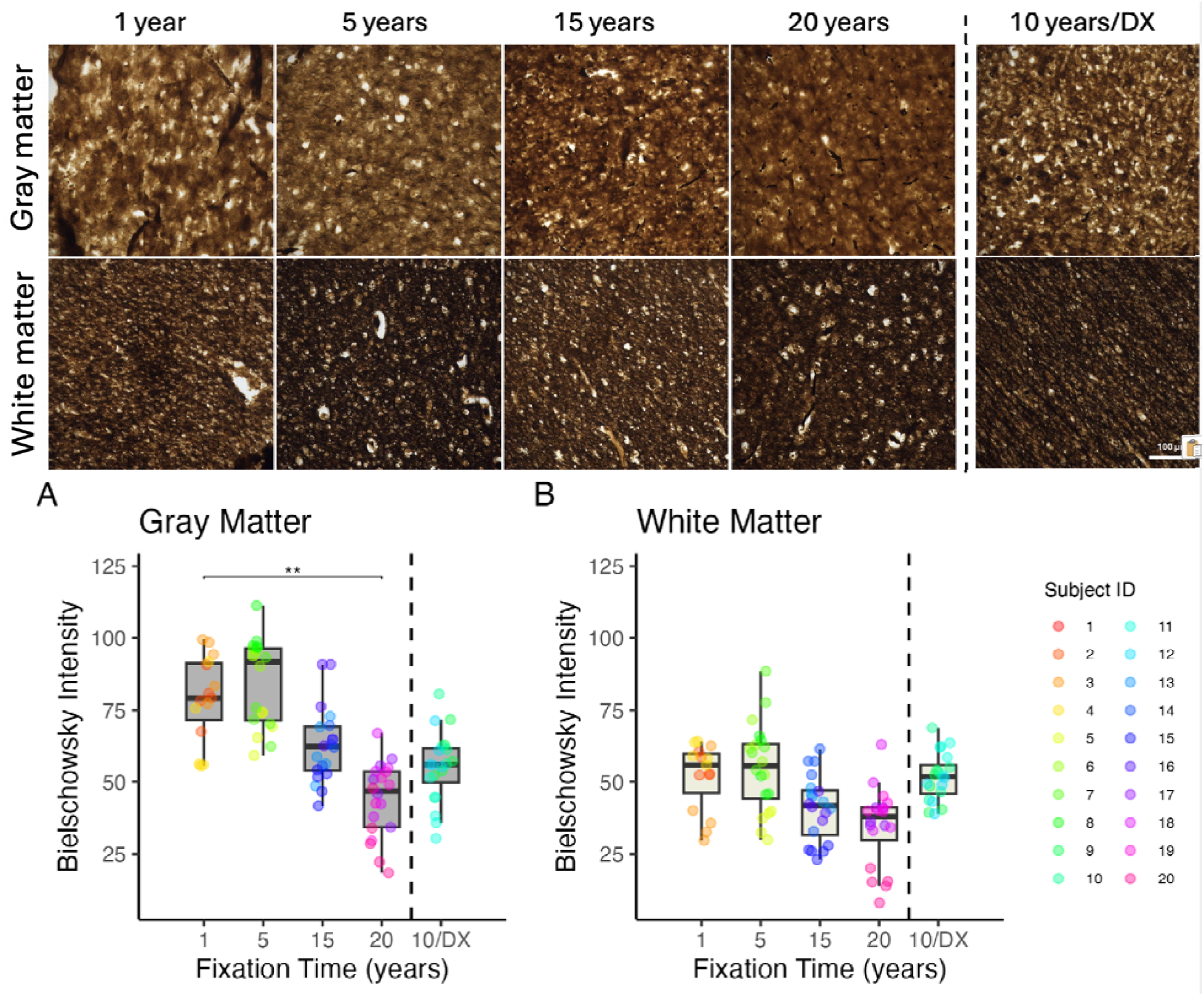
Bielschowsky silver staining in the gray and white matter (20X) of the PFC sections of human brains fixed for 1, 5, 10, 15, and 20 years. A) Mean staining intensity of the gray matter in the five groups. B) Mean staining intensity of the white matter in the five groups. Mean staining intensity was measured based on RGB intensity values. DX=Diagnosis; ID=Identification.

## 4. Discussion

In the present study, we systematically evaluated the impact of prolonged fixation on antibodies targeting various biomarkers on the PFC of human brains fixed for 1, 5, 10, 15, and 20 years. We found that prolonged fixation resulted in a reduction of the IHC staining intensity for Ferritin, Vimentin and Collagen-IV, as well as HC staining intensity reduction for BSS. Given that when looking at the micrographs, the cells can still be observed even if they are lightly stained in long-term fixed samples, this should not affect cell density estimations. Furthermore, we found no association between increased fixation and the IHC staining intensity of CD68, PLP and Claudin-5, nor between increased fixation and the HC staining intensity of MTS, indicating that their staining intensity remains mostly stable during fixation. Taken together, these data indicate that in human brains, prolonged fixation induces differential effects on commonly used biomarkers for CVD. Given that the biomarkers examined in this study are commonly used to investigate the pathogenesis of CVD and NDDs such as AD, our findings suggest that neuropathological assessments need to consider fixation length as a factor in their interpretations of staining intensity analyses.

Due to the protein and RNA crosslinking caused by NBF [37], it is generally believed that prolonged fixation with NBF produces IHC antigenicity reduction in human brain samples [13]. Prior studies on human brains showed a negative correlation between NeuN intensity and fixation times ranging from immediate up to 3 years [32] or from fixation durations spanning 24 hours, 4 months, and 10 years [30]. Animal studies also supported a similar decline in IHC staining by prolonged fixation. For example, NeuN levels were reduced in the swine brain fixed for 2 months, which completely disappeared after 3-months of fixation [38]. Similarly, prolonged fixation decreased IHC staining of biomarkers of neurogenesis in adult mice [39]. Consistent with these studies, we previously showed a negative correlation between IHC staining intensity of NeuN and Iba1 on human brain sections and fixation years [16]. We therefore expected that fixation duration could impact other markers that are commonly used in neuropathology or for studying vascular disease burden in the brain. However, our prior study showed that prolonged fixation enhanced the IHC staining of the astrocyte marker GFAP in human brains [16]. While NBF-induced protein crosslinking is often considered the main cause for the negative effect of fixation on IHC staining by masking out epitopes [13, 40], crosslinking can also be responsible for increased staining intensity and/or antigenicity detection. For example, we reasoned that more GFAP epitopes could be exposed by breaking protein crosslinking during antigen retrieval [16]. It is still unclear why some specific antigens (i.e., according to their specific biochemical protein folding) are differently affected by fixation. Taken together, these studies and ours suggest that prolonged fixation may have differential effects on IHC staining quality for some but not all target proteins. Therefore, further studies are required to provide additional evidence for these interpretations.

Since our previous study showed negative correlations between Iba1 staining intensity and fixation times [16], we expected that another microglia marker (CD68) would also be impacted by fixation times. The activated microglia marker CD68 is a more active player than other general microglia markers (such as Iba1) in neuropathology of AD brains [41]. CD68 is a widely used marker for identifying and visualizing microglia, particularly when they are in an activated state, as macrophages [42], and has been used extensively to identify inflammatory processes in neurodegenerative diseases [43–45]. Furthermore, microglia are known markers of neuroinflammation since their activation is implicated in CVD lesions, especially in WMH [46]. A prior qualitative study showed that CD68 immunoreactive activated microglia were observed in the frontal lobes of human brains fixed from < 1 year up to 20 years. While this is consistent with our current results, this study did not report any quantitative evidence on the correlation between CD68 IHC staining intensity and the fixation length [29]. Another study used CD68 IHC staining to assess Aβ immunized AD cases vs non-immunized AD cases [47]. However, the potential impact of the fixation delay in the quantitative analysis of the pathology associated with this biomarker was not examined. Even though not statistically significant, our data shows a declining trend in staining intensity with fixation years, suggesting that while CD68 markers may be used in long-term fixed samples, fixation times should be considered in the analyses.

PLP is known as the human proteolipid protein, a biomarker for oligodendrocytic myelin, a major structural component of myelin, helping to compact and stabilize the myelin sheath, which is vital for efficient nerve impulse transmission [48–50]. PLP IHC is widely applied for the neuropathological evaluation of demyelinating disorders, including multiple sclerosis [48, 49] and Pelizaeus-Merzbacher disease [51]. Furthermore, in the WM of AD brains, PLP decreases when the levels of amyloid-β increase [52], suggesting that abnormalities of myelin and oligodendrocytes are linked to AD pathology [53]. However, our 10-years / DX group did not show any significant decrease of PLP staining intensity in the WM. Overall, our results showed no significant changes in the staining intensity of PLP in brains fixed for decades. Therefore, given that our previous study in the same specimens showed a reduction in the staining intensity of Luxol fast blue which also stains myelin sheath [16], we recommend the use of PLP instead of Luxol fast blue in studies examining long-term fixed tissues.

Our ferritin marker stained iron deposits as well as microglia, since macrophages are known to perform phagocytosis of iron when neuroinflammation occurs in NDDs [54, 55]. Ferritin is a widely used biomarker to assess the extent of CVD lesions in stroke [56] and cerebral amyloid angiopathy [57–59], as well as normal iron accumulation in aging [60] or in NDDs [61–63]. IHC using ferritin or other iron markers are also essential to validate ex vivo MRI or quantitative susceptibility maps [64, 65]. However, to our knowledge, our study is the first to quantitatively evaluate the impact of fixation on the immunoreactivity of ferritin antibodies in postmortem human brains. Our results showed a significantly reduced staining intensity when brains were fixed for long periods. This should be considered in ex vivo studies that examine iron deposits using histopathology, although current studies mainly use Perl’s stain [8, 57, 66–68].

Claudin-5 is an essential protein for sealing the intercellular space between adjacent endothelial cells in the BBB (i.e., a biomarker for tight-junctions), primarily used to assess the integrity of the BBB and its potential disruption [69] which is involved in aging and AD [70]. It is also investigated as a biomarker for various other neurological and psychiatric disorders [71]. We found that Claudin-5 staining intensity remained stable during fixation, suggesting that it is a reliable marker for BBB assessment.

Vimentin is a marker for intermediate filament III, present in ependymal cells as well as in the cytoskeleton of astrocytes [72, 73]. It is also used as a marker for endothelial cells, particularly those lining blood vessels [74], and is used as a marker of BBB disruptions [75], reflecting its significance for CVD lesions studies. Furthermore, vimentin is closely associated with AD due to its role in Aß aggregation and deposition [76, 77]. Our current study showed that vimentin not only labeled blood vessels, but also activated astrocytes, components of the BBB, which is consistent with previous studies investigating vimentin co-localization with other astrocytes markers such as GFAP [78]. The latter study used postmortem human brain blocks of 1 cm^3^ that were fixed immediately in NBF after brain extraction. They showed very high staining quality of astrocytes and blood vessels [78], while other studies showed no changes in vimentin staining intensity in dog brains fixed for 7 weeks [26, 31]. This is also in agreement with our current study, where the 1-year group showed very high staining intensity of both cell-types, but that there is a significant decline in vimentin staining when brains are fixed for longer durations.

Collagen is a ubiquitous protein of the extracellular matrix of various connective tissues, with type I being the most widespread [79]. Collagen IV is a crucial component of blood vessel walls, contributing to their structural integrity and function [80], and its presence and turnover are important indicators of vascular health and BBB integrity, with imbalances potentially leading to various conditions [81]. Collagen fibers in the central nervous system contribute to NDDs, including AD [82]. This biomarker has been assessed using IHC in small vessel disease [83], cognitive impairment [84] and vascular dementia [85]. Our current findings also suggest that glial cell bodies are stained by Collagen-IV markers alongside blood vessels, representing the major components of the BBB, reflecting its significance as a marker of CVD. Previous studies have also investigated collagen as a potential marker for forensic evaluation, since the cellular matrix resists degradation during longer PMIs compared to the cellular components [79]. However, we showed that staining intensity of collagen is reduced in brains fixed for decades. These discrepancies in collagen IHC quality may be due to i) differences between Collagen-I and Collagen-IV studies; ii) tissue types (brain vs other forensic tissues); and iii) Collagen inside blood vessel walls vs the extracellular matrix. These data suggest that further studies should include fixation length as a covariate when investigating collagen fibers in the brain.

BSS is a well-known stain used for differentiating between NDDs through selective staining of neuritic plaques, neurofibrillary tangles, and their axonal prolongations with silver deposits, thereby visualizing the two characteristic pathological hallmarks of AD [10, 86–95]. Overall, BSS staining intensity decreased with fixation length reaching 20 years, since it is a silver revelation technique in which the tissue background may be affected by interaction with chemicals [15]. This suggests that, even though HC methods are not antigenicity-dependent (and therefore, not crosslinking dependent) compared to IHC methods, they can still be affected by prolonged chemical fixation, which should be taken into account when using these stains in future studies.

Finally, MTS is a common histopathological stain used to identify collagen and fibrin accumulation [96], as well as the extracellular matrix in brain tissue for BBB health and perivascular spaces (common markers of CVD lesions) [97]. MTS also stains myelin fibers within the extracellular matrix, allowing for the identification of CVD lesions, including WMHs [8, 98–100]. It is commonly used to visualize stenosis and amyloid-β and tau accumulation within blood vessels, which are characteristic of cerebral amyloid angiopathy in NDD patients or animal models [89]. Here, we found no significant differences in the staining intensity of MTS in groups fixed for decades, reflecting its robustness for use in multiple studies that employ brains fixed for years in brain banks.

At the time of our tissue request, our brain bank did not have any healthy-aging control PFC blocks available for the 10-years group (from the suicide brain bank). Instead of excluding this time frame from our study, we requested tissues with a diagnosis of CVD/NDDs for this group. This is a limitation also considering that the characteristics of the 10-years group were different from the others, especially since they were significantly older. However, including the 10-year/DX samples also allowed us to examine whether these markers could reliably detect the expected pathologies in a population of interest as we found differences in the staining intensities of markers that may be related to the underlying pathology of this group. For example, CD68 showed a wide range of staining intensities, from low values (3.68 in GM and 9.93 in WM) due to the decrease in staining intensity after 10 years of fixation, to higher values (20.46 in GM and 40.21 in WM), likely due to the increase of microglial activation resulting from neuroinflammation and neurodegeneration [43–45]. The highest values were obtained in Specimen 12, which was diagnosed with co-pathologies of AD, LBD, and CVD, suggesting increased microglial activation when more pathologies are involved, resulting in higher neuroinflammation [46]. Claudin-5 also showed significantly higher intensity values in the WM regions than the other groups. This may be because Claudin-5 is a tight junction protein involved in the BBB and may be dysregulated in AD [69, 70]. We expected to find differences in BSS since it is used as a marker of neuritic plaques and neurofibrillary tangles, especially in AD [10, 86–95]. Indeed, the presence of some neuritic plaques and neurofibrillary tangles were found in three out of four cases of the 10-years group. However, all of these differences in the 10-years/DX group did not reach significance after a FDR correction, potentially because this group was older, which results in a high collinearity between age and group variables, while our small sample size can limit our ability to detect significant differences.

Indeed, we acknowledge that we used a small sample to evaluate the fixation changes, which limited the power of our statistical analysis. However, this might be counterbalanced by the inclusion of a wide range of fixation times. This better reflects the actual samples that brain banks can provide, in which some blocks of tissue are taken from brains preserved in formaldehyde often for decades. Furthermore, as mentioned previously, changes observed in our 10-years group may have been impacted by presence of pathology and their difference in age. We did not assess the same cases longitudinally, which would have been ideal for evaluating fixation changes. However, this was not feasible for practical reasons, as tissue availability from the same specimens is limited and longitudinal sampling within our timeframe of interest was not feasible. Nevertheless, we consistently selected the same region in all cases and matched the cases as closely as possible for age and sex. Since age and sex had no statistically significant effects on CVD markers and that these can be detected both in normal aging and in NDDs, we believe the impact of these covariates on our findings was minimal. Another limitation of our study is that we did not optimize the protocols for each group (for example, antibody concentration or DAB incubation could have been increased in the groups fixed for longer durations). However, our goal was to assess the impact of the fixation length on the IHC staining quality, and modifications of the protocols between the fixation groups would have confounded comparisons. Finally, while we focused on CVD markers, future studies should include additional neurodegenerative markers, such as amyloid-β, phosphorylated tau, TDP-43 or α-synuclein, especially since they are widely used for neuropathology assessment of NDDs and dementia [77, 89, 101–104]. These markers were previously assessed according to fixation ranging from 0-14 years, where these studies also suggested differential effects of fixation lengths on different markers [105]. These markers would therefore be of interest in future studies.

In this study, IHC targeting six biomarkers and two HC protocols widely used in studying NDDs were examined in postmortem human brains fixed for 1, 5, 10, 15 or 20 years. Taken together, we conclude that prolonged fixation can produce negative effects on the IHC or HC staining intensities for human brain sections, depending on the biomarkers. In a nutshell, examining the effects of prolonged fixation on the IHC and HC staining quality of these biomarkers in human brains is necessary to enable accurate interpretation of subsequent experimental investigation of their related pathologies. Our findings suggest that studies using postmortem human brains to assess different biomarkers in CVD, NDD, or normal aging should take the fixation time into account as a covariate or use specimens fixed for the same durations for all analyses. A good knowledge of how antigens are affected by fixation is required before launching group comparison quantification studies, especially when measuring staining intensities.

## Funding

The current study was supported by research funds from Natural Sciences and Engineering Research Council of Canada (NSERC, Ref # RGPIN-2023-04038), Fonds de Recherche du Québec - Santé (FRQS, Ref # CB - 330750) and Canadian Institutes of Health Research (CIHR, Ref # 19130) awarded to Dr. Mahsa Dadar as well as funding from the Fonds de recherche du Québec—Santé (FRQS https://doi.org/10.69777/320107), Natural Sciences and Engineering Research Council of Canada (NSERC, Ref # RGPIN-2023-04218), and Canadian Institutes of Health Research (CIHR, PJT-195671) awarded to Dr. Yashar Zeighami. The DBB is supported by a Platform Support Grant from Brain Canada awarded to Gustavo Turecki and Naguib Mechawar.

## Competing interests

The authors report no competing interests with any parties.

## Notes

### Competing Interest Statement

The authors have declared no competing interest.

### Summary of Updates

Title changed. We added some co-authors.

## References

1. Cortes-Canteli, M. and C. Iadecola, Alzheimer’s Disease and Vascular Aging: JACC Focus Seminar. J Am Coll Cardiol, 2020. 75(8): p. 942–951.

2. Sabbatinelli, J., et al., Connecting vascular aging and frailty in Alzheimer’s disease. Mech Ageing Dev, 2021. 195: p. 111444.

3. Arvanitakis, Z., et al., Relation of cerebral vessel disease to Alzheimer’s disease dementia and cognitive function in elderly people: a cross-sectional study. Lancet Neurol, 2016. 15(9): p. 934–943.

4. Dadar, M., et al., White matter hyperintensity distribution differences in aging and neurodegenerative disease cohorts. Neuroimage Clin, 2022. 36: p. 103204.

5. Wardlaw, J.M., M.C. Valdés Hernández, and S. Muñoz-Maniega, What are white matter hyperintensities made of? Relevance to vascular cognitive impairment. J Am Heart Assoc, 2015. 4(6): p. 001140.

6. Cardoso, J.R., et al., What is gold standard and what is ground truth? Dental Press J Orthod, 2014. 19(5): p. 27–30.

7. Durand-Martel, P., et al., Autopsy as gold standard in FDG-PET studies in dementia. Can J Neurol Sci, 2010. 37(3): p. 336–42.

8. Humphreys, C.A., et al., A protocol for precise comparisons of small vessel disease lesions between ex vivo magnetic resonance imaging and histopathology. Int J Stroke, 2019. 14(3): p. 310–320.

9. Hoffmann, A., et al., Validation of in vivo magnetic resonance imaging blood-brain barrier permeability measurements by comparison with gold standard histology. Stroke, 2011. 42(7): p. 2054–60.

10. Kovacs, G.G. and H. Budka, Current concepts of neuropathological diagnostics in practice: neurodegenerative diseases. Clin Neuropathol, 2010. 29(5): p. 271–88.

11. Sukswai, N. and J.D. Khoury, Immunohistochemistry Innovations for Diagnosis and Tissue-Based Biomarker Detection. Curr Hematol Malig Rep, 2019. 14(5): p. 368–375.

12. Adickes, E.D., R.D. Folkerth, and K.L. Sims, Use of perfusion fixation for improved neuropathologic examination. Arch Pathol Lab Med, 1997. 121(11): p. 1199–206.

13. Stumptner, C., et al., The impact of crosslinking and non-crosslinking fixatives on antigen retrieval and immunohistochemistry. N Biotechnol, 2019. 52: p. 69–83.

14. Frigon, E.M., et al., Antigenicity is preserved with fixative solutions used in human gross anatomy: A mice brain immunohistochemistry study. Front Neuroanat, 2022. 16.

15. Frigon, E.M., et al., Comparison of histological procedures and antigenicity of human post-mortem brains fixed with solutions used in gross anatomy laboratories. Front Neuroanat, 2024. 18: p. 1372953.

16. Ma, W., et al., Differential effects of prolonged post-fixation on immunohistochemical and histochemical staining for postmortem human brains. Front Neuroanat, 2024. Volume 18 - 2024.

17. Hade, A.C., et al., A cost-effective and efficient ex vivo, ex situ human whole brain perfusion protocol for immunohistochemistry. J Neurosci Methods, 2024. 404: p. 110059.

18. Beach, T.G., et al., Arizona Study of Aging and Neurodegenerative Disorders and Brain and Body Donation Program. Neuropathology, 2015. 35(4): p. 354–89.

19. Vonsattel, J.P., et al., Twenty-first century brain banking: practical prerequisites and lessons from the past: the experience of New York Brain Bank, Taub Institute, Columbia University. Cell Tissue Bank, 2008. 9(3): p. 247–58.

20. McFadden, W.C., et al., Perfusion fixation in brain banking: a systematic review. Acta Neuropathol Commun, 2019. 7(1): p. 146.

21. Weisbecker, V., Distortion in formalin-fixed brains: Using geometric morphometrics to quantify the worst-case scenario in mice. Brain structure & function, 2011. 217: p. 677–85.

22. Helander, K.G., Kinetic studies of formaldehyde binding in tissue. Biotech Histochem, 1994. 69(3): p. 177–9.

23. Thavarajah, R., et al., Chemical and physical basics of routine formaldehyde fixation. J Oral Maxillofac Pathol, 2012. 16(3): p. 400–5.

24. Arnold, M.M., et al., Effects of fixation and tissue processing on immunohistochemical demonstration of specific antigens. Biotech Histochem, 1996. 71(5): p. 224–30.

25. Jiao, Y., et al., A simple and sensitive antigen retrieval method for free-floating and slide-mounted tissue sections. J Neurosci Methods, 1999. 93(2): p. 149–62.

26. Ramos-Vara, J.A. and M.E. Beissenherz, Optimization of immunohistochemical methods using two different antigen retrieval methods on formalin-fixed paraffin-embedded tissues: experience with 63 markers. J Vet Diagn Invest, 2000. 12(4): p. 307–11.

27. Shi, S.R., R.J. Cote, and C.R. Taylor, Antigen retrieval immunohistochemistry used for routinely processed celloidin-embedded human temporal bone sections: standardization and development. Auris Nasus Larynx, 1998. 25(4): p. 425–43.

28. Shi, S.R., Y. Shi, and C.R. Taylor, Antigen retrieval immunohistochemistry: review and future prospects in research and diagnosis over two decades. J Histochem Cytochem, 2011. 59(1): p. 13–32.

29. Alrafiah, A. and R. Alshali, The effect of prolonged formalin fixation on the staining characteristics of archival human brain tissue. Folia Morphol (Warsz), 2019. 78(2): p. 230–236.

30. Lyck, L., et al., Immunohistochemical markers for quantitative studies of neurons and glia in human neocortex. J Histochem Cytochem, 2008. 56(3): p. 201–21.

31. Webster, J.D., et al., Effects of prolonged formalin fixation on diagnostic immunohistochemistry in domestic animals. J Histochem Cytochem, 2009. 57(8): p. 753–61.

32. Wu, X., et al., The effect of prolonged formalin fixation on the expression of proteins in human brain tissues. Acta Histochem, 2022. 124(4): p. 151879.

33. Tovi, M. and A. Ericsson, Measurements of T1 and T2 over time in formalin-fixed human whole-brain specimens. Acta Radiol, 1992. 33(5): p. 400–4.

34. Dadar, M., et al., The Douglas-Bell Canada Brain Bank Post-mortem Brain Imaging Protocol. Aperture Neuro, 2024. 4.

35. Shu, S.Y., G. Ju, and L.Z. Fan, The glucose oxidase-DAB-nickel method in peroxidase histochemistry of the nervous system. Neurosci Lett, 1988. 85(2): p. 169–71.

36. Benjamini, Y. and Y. Hochberg, Controlling the False Discovery Rate: A Practical and Powerful Approach to Multiple Testing. Journal of the Royal Statistical Society: Series B (Methodological), 1995. 57(1): p. 289–300.

37. Kiernan, J.A., Formaldehyde, Formalin, Paraformaldehyde And Glutaraldehyde: What They Are And What They Do. Microscopy Today, 2000. 8(1): p. 8–13.

38. Lundström, Y., et al., Detection of Changes in Immunohistochemical Stains Caused by Postmortem Delay and Fixation Time. Appl Immunohistochem Mol Morphol, 2019. 27(3): p. 238–245.

39. Gallardo-Caballero, M., et al., Prolonged fixation and post-mortem delay impede the study of adult neurogenesis in mice. Commun Biol, 2023. 6(1): p. 978.

40. Hoffman, E.A., et al., Formaldehyde crosslinking: a tool for the study of chromatin complexes. J Biol Chem, 2015. 290(44): p. 26404–11.

41. Hopperton, K.E., et al., Markers of microglia in post-mortem brain samples from patients with Alzheimer’s disease: a systematic review. Mol Psychiatry, 2018. 23(2): p. 177–198.

42. Zigmond, R.E. and F.D. Echevarria, Macrophage biology in the peripheral nervous system after injury. Prog Neurobiol, 2019. 173: p. 102–121.

43. Farso, M., et al., The immune marker CD68 correlates with cognitive impairment in normally aged rats. Neurobiology of Aging, 2013. 34(8): p. 1971–1976.

44. Minett, T., et al., Microglial immunophenotype in dementia with Alzheimer’s pathology. J Neuroinflammation, 2016. 13(1): p. 135.

45. Wetering, J.V., et al., Neuroinflammation is associated with Alzheimer’s disease co-pathology in dementia with Lewy bodies. Acta Neuropathol Commun, 2024. 12(1): p. 73.

46. Makkinejad, N., et al., Neuropathological Correlates of White Matter Hyperintensities in Cerebral Amyloid Angiopathy. J Am Heart Assoc, 2024. 13(22): p. e035744.

47. Zotova, E., et al., Microglial alterations in human Alzheimer’s disease following Aβ42 immunization. Neuropathol Appl Neurobiol, 2011. 37(5): p. 513–24.

48. Greer, J.M. and M.P. Pender, Myelin proteolipid protein: An effective autoantigen and target of autoimmunity in multiple sclerosis. Journal of Autoimmunity, 2008. 31(3): p. 281–287.

49. Greer, J.M., E. Trifilieff, and M.P. Pender, Correlation Between Anti-Myelin Proteolipid Protein (PLP) Antibodies and Disease Severity in Multiple Sclerosis Patients With PLP Response-Permissive HLA Types. Front Immunol, 2020. 11: p. 1891.

50. Knapp, P.E., Proteolipid protein: is it more than just a structural component of myelin? Dev Neurosci, 1996. 18(4): p. 297–308.

51. Simons, M., et al., Overexpression of the myelin proteolipid protein leads to accumulation of cholesterol and proteolipid protein in endosomes/lysosomes: implications for Pelizaeus-Merzbacher disease. J Cell Biol, 2002. 157(2): p. 327–36.

52. Roher, A.E., et al., Increased A beta peptides and reduced cholesterol and myelin proteins characterize white matter degeneration in Alzheimer’s disease. Biochemistry, 2002. 41(37): p. 11080–90.

53. Nasrabady, S.E., et al., White matter changes in Alzheimer’s disease: a focus on myelin and oligodendrocytes. Acta Neuropathol Commun, 2018. 6(1): p. 22.

54. Andersen, H.H., K.B. Johnsen, and T. Moos, Iron deposits in the chronically inflamed central nervous system and contributes to neurodegeneration. Cell Mol Life Sci, 2014. 71(9): p. 1607–22.

55. Simmons, D.A., et al., Ferritin accumulation in dystrophic microglia is an early event in the development of Huntington’s disease. Glia, 2007. 55(10): p. 1074–84.

56. Justicia, C., P. Ramos-Cabrer, and M. Hoehn, MRI detection of secondary damage after stroke: chronic iron accumulation in the thalamus of the rat brain. Stroke, 2008. 39(5): p. 1541–7.

57. Guidoux, C., et al., Amyloid Angiopathy in Brain Hemorrhage: A Postmortem Neuropathological-Magnetic Resonance Imaging Study. Cerebrovasc Dis, 2018. 45(3-4): p. 124–131.

58. Schrag, M., et al., Effect of cerebral amyloid angiopathy on brain iron, copper, and zinc in Alzheimer’s disease. J Alzheimers Dis, 2011. 24(1): p. 137–49.

59. Sharma, B., et al., Brain iron content in cerebral amyloid angiopathy using quantitative susceptibility mapping. Front Neurosci, 2023. 17: p. 1139988.

60. Connor, J.R., et al., Cellular distribution of transferrin, ferritin, and iron in normal and aged human brains. J Neurosci Res, 1990. 27(4): p. 595–611.

61. Daglas, M. and P.A. Adlard, The Involvement of Iron in Traumatic Brain Injury and Neurodegenerative Disease. Front Neurosci, 2018. 12: p. 981.

62. Long, H., et al., Iron homeostasis imbalance and ferroptosis in brain diseases. MedComm (2020), 2023. 4(4): p. e298.

63. van Duijn, S., et al., Comparison of histological techniques to visualize iron in paraffin-embedded brain tissue of patients with Alzheimer’s disease. J Histochem Cytochem, 2013. 61(11): p. 785–92.

64. Antharam, V., et al., High field magnetic resonance microscopy of the human hippocampus in Alzheimer’s disease: quantitative imaging and correlation with iron. Neuroimage, 2012. 59(2): p. 1249–60.

65. De Barros, A., et al., Matching ex vivo MRI With Iron Histology: Pearls and Pitfalls. Frontiers in neuroanatomy, 2019. 13: p. 68–68.

66. Nag, S., et al., Ex vivo MRI facilitates localization of cerebral microbleeds of different ages during neuropathology assessment. Free Neuropathol, 2021. 2: p. 35.

67. Schrag, M., et al., Correlation of hypointensities in susceptibility-weighted images to tissue histology in dementia patients with cerebral amyloid angiopathy: a postmortem MRI study. Acta Neuropathol, 2010. 119(3): p. 291–302.

68. Tatsumi, S., M. Shinohara, and T. Yamamoto, Direct comparison of histology of microbleeds with postmortem MR images: a case report. Cerebrovasc Dis, 2008. 26(2): p. 142–6.

69. Lv, J., et al., Focusing on claudin-5: A promising candidate in the regulation of BBB to treat ischemic stroke. Progress in Neurobiology, 2018. 161: p. 79–96.

70. Tachibana, K., et al., Association of Plasma Claudin-5 with Age and Alzheimer Disease. Int J Mol Sci, 2024. 25(3).

71. Greene, C., N. Hanley, and M. Campbell, Claudin-5: gatekeeper of neurological function. Fluids Barriers CNS, 2019. 16(1): p. 3.

72. Chiu, F.C., W.T. Norton, and K.L. Fields, The cytoskeleton of primary astrocytes in culture contains actin, glial fibrillary acidic protein, and the fibroblast-type filament protein, vimentin. J Neurochem, 1981. 37(1): p. 147–55.

73. Schnitzer, J., W.W. Franke, and M. Schachner, Immunocytochemical demonstration of vimentin in astrocytes and ependymal cells of developing and adult mouse nervous system. J Cell Biol, 1981. 90(2): p. 435–47.

74. Kidd, M.E., D.K. Shumaker, and K.M. Ridge, The role of vimentin intermediate filaments in the progression of lung cancer. Am J Respir Cell Mol Biol, 2014. 50(1): p. 1–6.

75. Bayir, E. and A. Sendemir, Role of Intermediate Filaments in Blood-Brain Barrier in Health and Disease. Cells, 2021. 10(6).

76. Chen, K.Z., et al., Vimentin as a potential target for diverse nervous system diseases. Neural Regen Res, 2023. 18(5): p. 969–975.

77. Zhang, L., et al., Vimentin Fragmentation and Its Role in Amyloid-Beta Plaque Deposition in Alzheimer’s Disease. Int J Mol Sci, 2025. 26(7).

78. O’Leary, L.A., et al., Characterization of Vimentin-Immunoreactive Astrocytes in the Human Brain. Front Neuroanat, 2020. 14: p. 31.

79. Mazzotti, M.C., et al., Determining the time of death by morphological and immunohistochemical evaluation of collagen fibers in postmortem gingival tissues. Leg Med (Tokyo), 2019. 39: p. 1–8.

80. Chello, M., et al., Analysis of collagen and elastin content in primary varicose veins. J Vasc Surg, 1994. 20(3): p. 490.

81. Kim, H.L., Arterial stiffness and hypertension. Clin Hypertens, 2023. 29(1): p. 31.

82. Wareham, L.K., et al., Collagen in the central nervous system: contributions to neurodegeneration and promise as a therapeutic target. Mol Neurodegener, 2024. 19(1): p. 11.

83. Kumar, A.A., et al., Vascular Collagen Type-IV in Hypertension and Cerebral Small Vessel Disease. Stroke, 2022. 53(12): p. 3696–3705.

84. Özkan, E., et al., Blood-brain barrier leakage and perivascular collagen accumulation precede microvessel rarefaction and memory impairment in a chronic hypertension animal model. Metab Brain Dis, 2021. 36(8): p. 2553–2566.

85. Denver, P., et al., A Novel Model of Mixed Vascular Dementia Incorporating Hypertension in a Rat Model of Alzheimer’s Disease. Front Physiol, 2019. 10: p. 1269.

86. Allsop, D., Introduction to Alzheimer’s disease. Methods Mol Med, 2000. 32: p. 1–21.

87. Elobeid, A., et al., Alzheimer’s disease-related plaques in nondemented subjects. Alzheimers Dement, 2014. 10(5): p. 522–9.

88. Fan, S., et al., Curcumin-loaded PLGA-PEG nanoparticles conjugated with B6 peptide for potential use in Alzheimer’s disease. Drug Deliv, 2018. 25(1): p. 1091–1102.

89. Klakotskaia, D., et al., Memory deficiency, cerebral amyloid angiopathy, and amyloid-β plaques in APP+PS1 double transgenic rat model of Alzheimer’s disease. PLoS One, 2018. 13(4): p. e0195469.

90. Love, S. and J.A. Nicoll, Comparison of modified Bielschowsky silver impregnation and anti-ubiquitin immunostaining of cortical and nigral Lewy bodies. Neuropathol Appl Neurobiol, 1992. 18(6): p. 585–92.

91. Stahnisch, F.W., Max Bielschowsky (1869-1940). J Neurol, 2015. 262(3): p. 792–4.

92. Switzer, R.C., 3rd, Application of silver degeneration stains for neurotoxicity testing. Toxicol Pathol, 2000. 28(1): p. 70–83.

93. Uchihara, T., Silver diagnosis in neuropathology: principles, practice and revised interpretation. Acta Neuropathol, 2007. 113(5): p. 483–99.

94. Wisniewski, H.M., G.Y. Wen, and K.S. Kim, Comparison of four staining methods on the detection of neuritic plaques. Acta Neuropathol, 1989. 78(1): p. 22–7.

95. Yamamoto, T. and A. Hirano, A comparative study of modified Bielschowsky, Bodian and thioflavin S stains on Alzheimer’s neurofibrillary tangles. Neuropathol Appl Neurobiol, 1986. 12(1): p. 3–9.

96. Islam, M.A. and S. Kumar, Masson’s Trichrome Staining Technique to Evaluate Tissue Fibrosis. Methods Mol Biol, 2026. 2983: p. 91–100.

97. Onoda, A., et al., A Novel Staining Method for Detection of Brain Perivascular Injuries Induced by Nanoparticle: Periodic Acid-Schiff and Immunohistochemical Double-Staining. Front Toxicol, 2022. 4: p. 825984.

98. Bearer, E.L., Exploring Vascular Contributions to Cognitive Impairment: Small-Vessel Disease of White Matter and Microplastics/Nanoplastics. Am J Pathol, 2025. 195(11): p. 2059–74.

99. Fazekas, F., et al., The morphologic correlate of incidental punctate white matter hyperintensities on MR images. AJNR Am J Neuroradiol, 1991. 12(5): p. 915–21.

100. Keith, J., et al., Collagenosis of the Deep Medullary Veins: An Underrecognized Pathologic Correlate of White Matter Hyperintensities and Periventricular Infarction? J Neuropathol Exp Neurol, 2017. 76(4): p. 299–312.

101. Baker-Nigh, A., et al., Neuronal amyloid-β accumulation within cholinergic basal forebrain in ageing and Alzheimer’s disease. Brain, 2015. 138(Pt 6): p. 1722–37.

102. Frigerio, I., et al., Amyloid-β, p-tau and reactive microglia are pathological correlates of MRI cortical atrophy in Alzheimer’s disease. Brain Commun, 2021. 3(4): p. fcab281.

103. Hampel, H., et al., The Amyloid-β Pathway in Alzheimer’s Disease. Mol Psychiatry, 2021. 26(10): p. 5481–5503.

104. Marsh, S.E. and M. Blurton-Jones, Examining the mechanisms that link β-amyloid and α-synuclein pathologies. Alzheimers Res Ther, 2012. 4(2): p. 11.

105. Pikkarainen, M., P. Martikainen, and I. Alafuzoff, The Effect of Prolonged Fixation Time on Immunohistochemical Staining of Common Neurodegenerative Disease Markers. Journal of Neuropathology & Experimental Neurology, 2010. 69(1): p. 40–52.

